# Genetic Mapping of Monocyte Fate Decisions Following Myocardial Infarction

**DOI:** 10.1101/2023.12.24.573263

**Authors:** Andrew L Koenig, Farid F Kadyrov, Junedh M Amrute, Steven Yang, Carla J Weinheimer, Jessica M Nigro, Attila Kovacs, Gabriella Smith, Kory J Lavine

## Abstract

Inflammation contributes to the pathogenesis of myocardial infarction and heart failure and represents a viable therapeutic target. Monocytes and their progeny are highly abundant and display incredible functional diversity, serving as key determinants of myocardial inflammation and tissue repair. Much remains to be learned regarding mechanisms and signaling events that instruct monocyte fate decisions. We devised a genetic lineage tracing strategy using *Ccr2^crERT2^Rosa26^LSL-tdTomato^* mice in combination with single cell RNA-sequencing to map the differentiation trajectories of monocytes that infiltrate the heart after reperfused myocardial infarction. Monocytes are recruited to the heart early after injury and give rise to transcriptionally distinct and spatially restricted macrophage and dendritic cell-like subsets that are specified prior to extravasation and chronically persist within the myocardium. Pseudotime analysis predicted two differentiation trajectories of monocyte-derived macrophages that are partitioned into the border and infarct zones, respectively. Among these trajectories, we show that macrophages expressing a type I IFN responsive signature are an intermediate population that gives rise to MHC-II^hi^ macrophages, are localized within the border zone, and promote myocardial protection. Collectively, these data uncover new complexities of monocyte differentiation in the infarcted heart and suggest that modulating monocyte fate decisions may have clinical implications.

## Introduction

Myocardial infarction (MI) remains a leading cause of death and major cause of heart failure worldwide (1, 2). It is well established that ischemic damage to the heart promotes myocardial inflammation by triggering recruitment of peripheral leukocytes that impart a second hit to the heart and contribute to additional cardiomyocyte loss, fibrosis, adverse left ventricular (LV) remodeling, and impaired cardiac function. Among recruited leukocytes, monocytes and macrophages represent the most abundant immune cells to accumulate within the infarcted and failing heart (3).

Within the past decade, it has been established that cardiac macrophages are functionally heterogeneous and can be broadly divided into two populations distinguished by expression of C-C chemokine receptor 2 (CCR2). CCR2^−^ macrophages are an embryonically derived, long-lived population that comprises the major macrophage subset in the healthy heart. CCR2^−^ macrophages are self-replenishing, express conserved markers of tissue resident macrophages, and function to orchestrate cardiac development, homeostasis, adaptation, and repair (4–9). In contrast, CCR2^+^ macrophages are an inflammatory population derived from recruited Ly6c^hi^ monocytes that accumulate within the injured and chronically failing heart. CCR2^+^ macrophages are associated with and causally linked to heart failure as they contribute to damaging inflammation, myocardial fibrosis, and adverse LV remodeling (6, 10–17).

Recent studies from our group have leveraged single cell transcriptomics in both mice and humans to demonstrate additional complexity within cardiac monocyte and macrophage populations (3, 18–22). These studies have uncovered incredible diversity amongst CCR2^+^ monocyte-derived macrophages that infiltrate the infarcted heart. While these findings have suggested that monocytes may differentiate into functional diverse sub-populations of CCR2^+^ macrophages, little is known regarding their spatial-temporal dynamics, trajectories, and mechanisms by which monocytes differentiate into these populations.

Here, we construct a map of monocyte differentiation trajectories in reperfused myocardial infarction (MI) using single cell RNA sequencing. We demonstrate that monocytes are recruited to the heart early after MI and give rise to transcriptionally distinct and spatially restricted macrophage and dendritic cell-like subsets that chronically persist within the myocardium. Monocyte-derived cells adopt macrophage and dendritic cell fates as they extravasate out of the vascular compartment and dynamically shift their phenotype over time. We leveraged Pseudotime analysis and spatial transcriptomics to predict spatially restricted monocyte differentiation trajectories. Finally, we provide experimental validation and show that macrophages expressing a type I IFN responsive signature are an intermediate population that gives rise to MHC-II^hi^ macrophages and that this trajectory confers protection following myocardial injury. These findings highlight unrecognized complexities of monocyte differentiation and suggest that modulating monocyte fate decisions may have clinical implications.

## Results

### Monocytes are recruited to the heart early after MI and their progeny chronically persist within the myocardium

To trace the fates of monocytes and their progeny after MI, we subjected *Ccr2^CreERT2^Rosa26^LSL-tdTomato^* mice to closed-chest ischemia reperfusion injury to model reperfused MI.(23–25) *Ccr2^CreERT2^* mice harbor a tamoxifen inducible CRE recombinase inserted into the 3’ UTR of the *Ccr2* locus. We optimized a tamoxifen induction strategy to exclusively label circulating monocytes and not cardiac macrophages. A single injection of tamoxifen (60mg/kg, IP) into naïve mice achieved selective tdTomato labeling of Ly6c^hi^ monocytes in the blood at >80% efficiency 24 hours after injection. Cardiac macrophages did not express tdTomato (**Fig. 1A**). We also did not detect significant tdTomato labeling of neutrophils or lymphocytes in the blood or their progenitors within the bone marrow compartment (**Suppl. Fig. 1**). As such, this strategy allows lineage tracing of infiltrating monocytes and their progeny independent of resident macrophages.

**Fig. 1.**
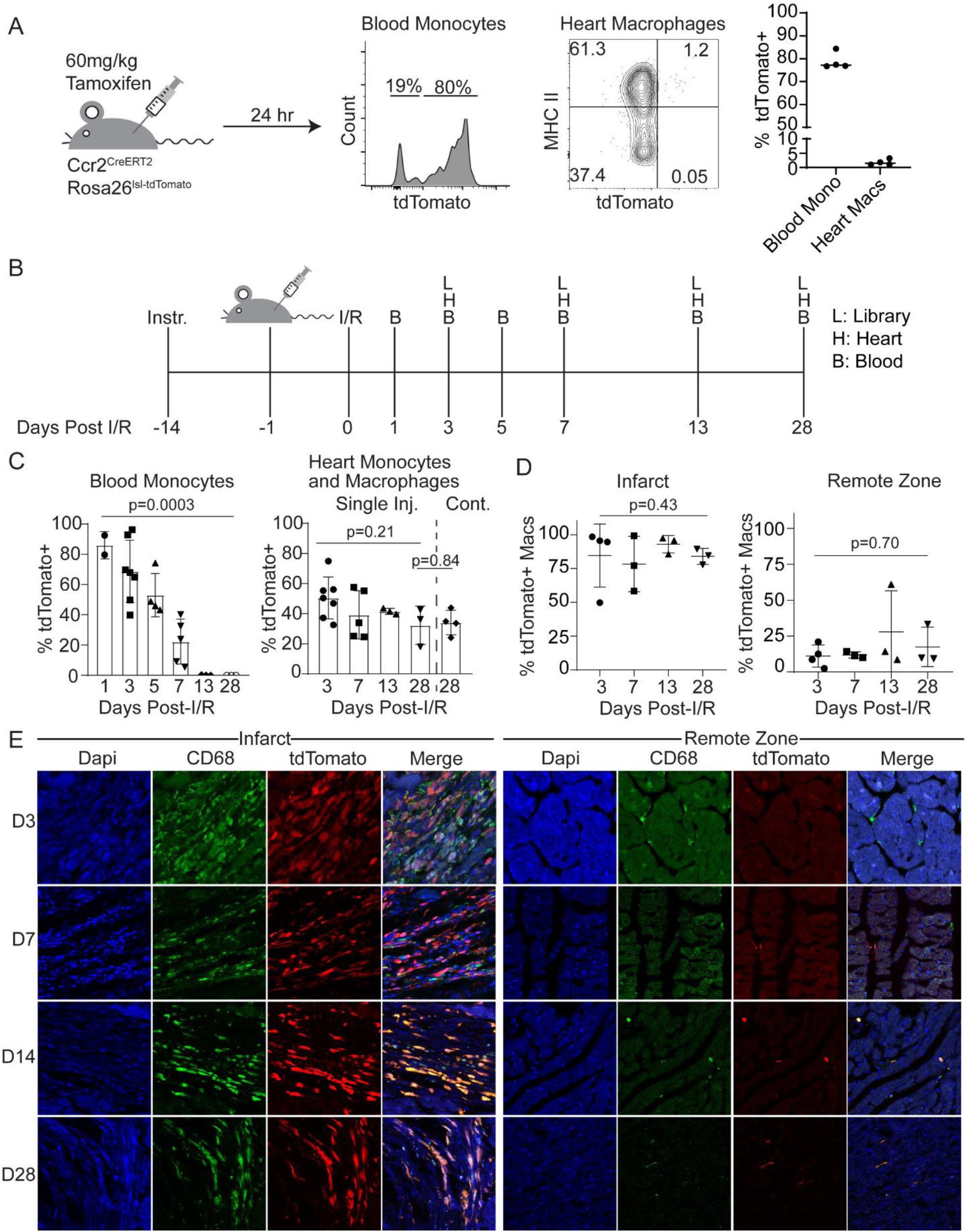
Recruited monocyte-derived populations persist in the heart after injury. **A**, Representative flow cytometry plots demonstrating that a single dose of tamoxifen administered to *Ccr2Cre^ERT2^Rosa26^tdTomato^* mice results in 80% labeling efficiency in monocytes in the blood, while resident cardiac macrophages are not labeled. **B**, Schematic of experimental strategy for monocyte lineage tracing after MI. **C**, Bar graphs demonstrating tdTomato labeling of blood monocytes (left) and cardiac monocyte and macrophages (right, single inj: single dose labeling, cont: continuous dose labeling). **D**, Quantification of tdTomato+ macrophages relative to total CD68 positive macrophages measured by IHC. **E**, Representative IHC images. Each data point represents an individual animal.

We then applied this labeling strategy to mice subjected to closed-chest ischemia reperfusion injury. Tamoxifen was injected 1 day prior to injury. Blood was collected from mice at days 1, 3, 5, 7, 13, and 28 after injury and hearts were collected at days 3, 7, 13, and 28 after injury (**Fig. 1B**). Flow cytometry analysis demonstrated >80% tdTomato labeling efficiency in Ly6c^hi^ monocytes in the blood 1 day post-MI. The percent of Ly6c^hi^ monocytes labeled progressively decreased throughout the first 7 days post-MI and was absent thereafter. Intriguingly, the progeny of monocytes that entered the heart during this labeling window persisted for at least 28 days post-MI. Continuous labeling of monocytes by tamoxifen injection (60mg/kg IP, every 3 days) did not demonstrate additional labeling of cardiac macrophages (**Fig. 1C**). Labeling of macrophages within the heart was validated at each time point by visualization of tdTomato and immunostaining for CD68 (**Fig. 1D-E**). These findings indicate that monocytes enter the heart predominately within the first 7 days post-MI and their progeny including monocyte-derived macrophages persist within the myocardium for at least 28 days post-MI.

### Macrophage states dynamically shift over time after injury

To delineate the temporal dynamics of recruited monocytes and their progeny, we performed single cell RNA-sequencing of monocytes and macrophages sorted from *Ccr2^CreERT2^Rosa26^LSL-tdTomato^* hearts collected at days 3, 7, 13, and 28 post-MI (**Fig. 1B, Suppl. Fig. 2A**). Libraries were constructed for each time point, sequenced, aligned using Cellranger, and then integrated, clustered, and analyzed using Seurat (26, 27). We identified 10 transcriptionally distinct cell clusters: monocytes, dendritic cells, and 8 macrophage states (**Fig. 2A**). We performed differential expression analysis and constructed z-score signatures to validate that the observed clusters were transcriptionally distinct (**Fig. B-C, Suppl. Fig. 3**). When cells were compared across time points, it was evident that macrophage states significantly shifted over time. At 3 days post-MI, the predominant populations within the heart consisted of monocytes, proliferating macrophages, *Arg1^+^* macrophages, type-1 interferon activated macrophages, and *Trem2^+^* macrophages. Through days 7 and 13 post-MI we observed decreases in frequency of monocytes, proliferating cells, *Arg1^+^* macrophages, and type-1 interferon activated macrophages. *Trem2^+^* macrophages and *Gdf15^+^* macrophages expanded during this time interval. By day 28 post MI, macrophages had transitioned to a predominantly MHCII^hi^ state (**Fig. 2D-E**).

**Fig. 2.**
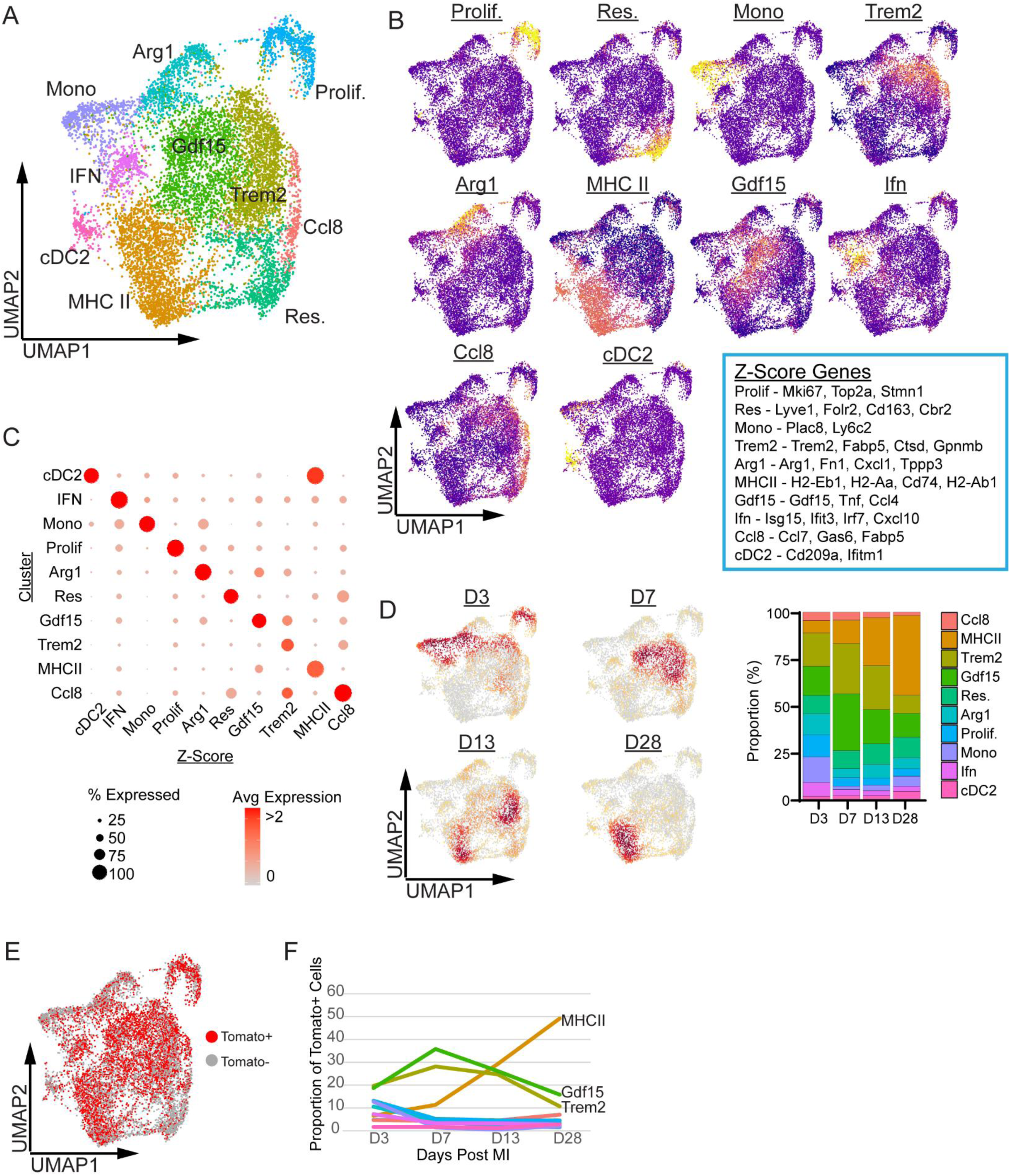
Recruited monocyte-derived populations are transcriptionally diverse and dynamically shift as a function of time after MI. **A**, Clustered UMAP projection of combined monocyte lineage tracing single cell libraries from each time point. **B**, Feature plots of calculated z-scores for each cluster (blue box: list of genes in each z-score). **C**, Dot plot of z-scores by cluster. **D**, Gaussian kernel plots indicating cell density at each time point after MI (left) and proportion of cells in each cluster split by time point (right). **E**, UMAP plot of Tomato+ cells. **F**, Line graph of proportion of tdTomato+ cells in each cluster across time points.

We additionally performed custom alignment of the libraries to the tdTomato coding sequence and observed that all identified clusters were labeled by expression of tdTomato. Interestingly, a proportion of macrophages within the resident cluster were also labeled by tdTomato indicating that recruited macrophages have the potential to adopt a resident-like fate. When comparing differential expression of tdTomato^−^ and tdTomato^+^ resident macrophages, we identified that tdTomato^+^ resident-like macrophages demonstrate reduced expression of the resident macrophage markers *Folr2* and *Lyve1* and enriched expression *Ccl2, Ccl7,* and *Ccl12* compared to tdTomato^−^ resident macrophages (**Fig. 2F, Suppl. Fig. 3**).

We next examined the dynamics of monocyte-derived cells as a function of time and found that tdTomato^+^ cells were substantially distributed across cell states at day 3 post-MI. At days 7 and 14 post-MI the number of tdTomato^+^ monocytes, *Arg1^+^* macrophages, type-1 interferon activated macrophages, and proliferating macrophages declined and a shift towards *Gdf15^+^* and *Trem2^+^* macrophages was evident. At day 28 post-MI the majority of tdTomato^+^ cells were MHCII^hi^ macrophages (**Fig. 2G**). Together, these results demonstrate that monocyte recruitment occurs within the first week after MI, their progeny persist within the heart at chronic stages after MI, and that monocytes display incredible plasticity differentiating into multiple transcriptionally distinct subsets that dynamically shift over time.

### Monocyte-derived macrophage and dendritic cell-like states exist outside of resident cardiac macrophages

We utilized *Cx3cr1^CreERT2^Rosa26^LSL-tdTomato^* mice to evaluate the contribution of resident macrophages to the diverse subsets of monocyte-derived cells that emerge following MI. Utilizing a pulse-chase strategy, we injected mice with tamoxifen (60mg/kg daily X5 days) 2 weeks prior to ischemia reperfusion injury (**Fig. 3A**). This strategy specifically labels long-lived resident macrophages with tdTomato. Monocytes are not labeled as they are replenished by unlabeled monocytes during the chase period (13). Macrophage and monocytes were sorted via FACS into tdTomato^+^ (resident) and tdTomato^−^ (recruited) libraries for single cell RNA sequencing (**Fig. 3A, Suppl. Fig. 2B**). We observed that resident macrophages clustered independently of recruited macrophages and displayed distinct transcriptional profiles. Differentially gene expression analysis demonstrated that resident and recruited macrophages express distinct transcriptional signatures. Many of the genes enriched in monocyte-derived macrophages were markers of the diverse populations seen in *Ccr2^CreERT2^* lineage tracing. Conversely, classical markers of tissue resident macrophages were enriched within the resident cardiac macrophage library (*Cd163, Mrc1, Cd207, Vsig4, Folr2, Cbr2*) (**Fig. 3B-C**). To validate that the resident macrophage lineage tracing labeling was efficient, we aligned the single cell libraries to the tdTomato coding sequence and found that tdTomato was exclusively expressed in cells found within the resident macrophage library (**Fig. 3D-E**). These data indicate that resident macrophages do not contribute to the diverse macrophage subsets marked by *Ccr2-CreERT2* lineage tracing, consistent with their origin from infiltrating monocytes.

**Fig. 3.**
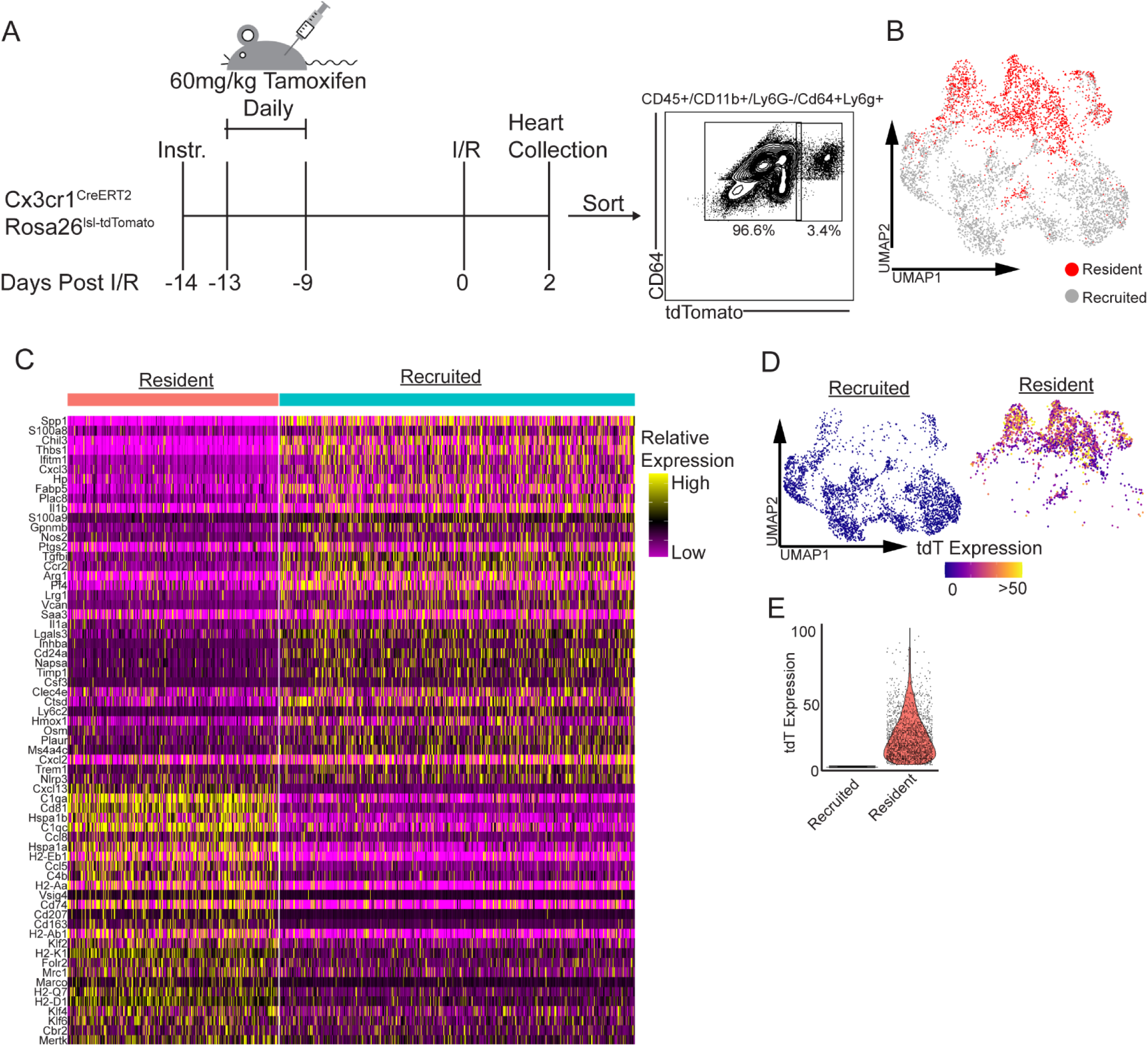
Monocyte-derived populations exist independent of cardiac resident macrophages. **A**, Schematic of experimental strategy for resident macrophage lineage tracing (left) and FACS gating strategy for constructing tdTomato+ (resident) and tdTomato-(recruited) libraries. **B**, Combined UMAP plot of resident and recruited libraries. Red indicates cells derived from cardiac resident macrophages. **C**, Heatmap of genes differentially expressed in cells derived from either cardiac resident macrophages or recruited monocytes. **D**, Feature plot of tdTomato expression split by library. **E**, Violin plot of tdTomato expression split by library.

### Macrophage states identified after myocardial infarction in mice are conserved in human heart failure

We utilized data from a prior study to determine if the observed monocyte, macrophage, and dendritic cell populations were conserved in the human heart. Transcription signatures for each mouse cluster were converted to their corresponding human homologues. Z-scores for these marker genes were calculated across human cardiac monocytes, macrophages, and dendritic cell clusters (**Fig. 4A-C**) (28). This analysis demonstrated that transcriptional signatures enriched in each mouse monocyte, macrophage, and dendritic cell cluster was conserved in corresponding populations of monocyte, macrophages, and dendritic cells found in the human heart indicating that mice may serve as an appropriate model to investigate human monocyte differentiation.

**Fig. 4.**
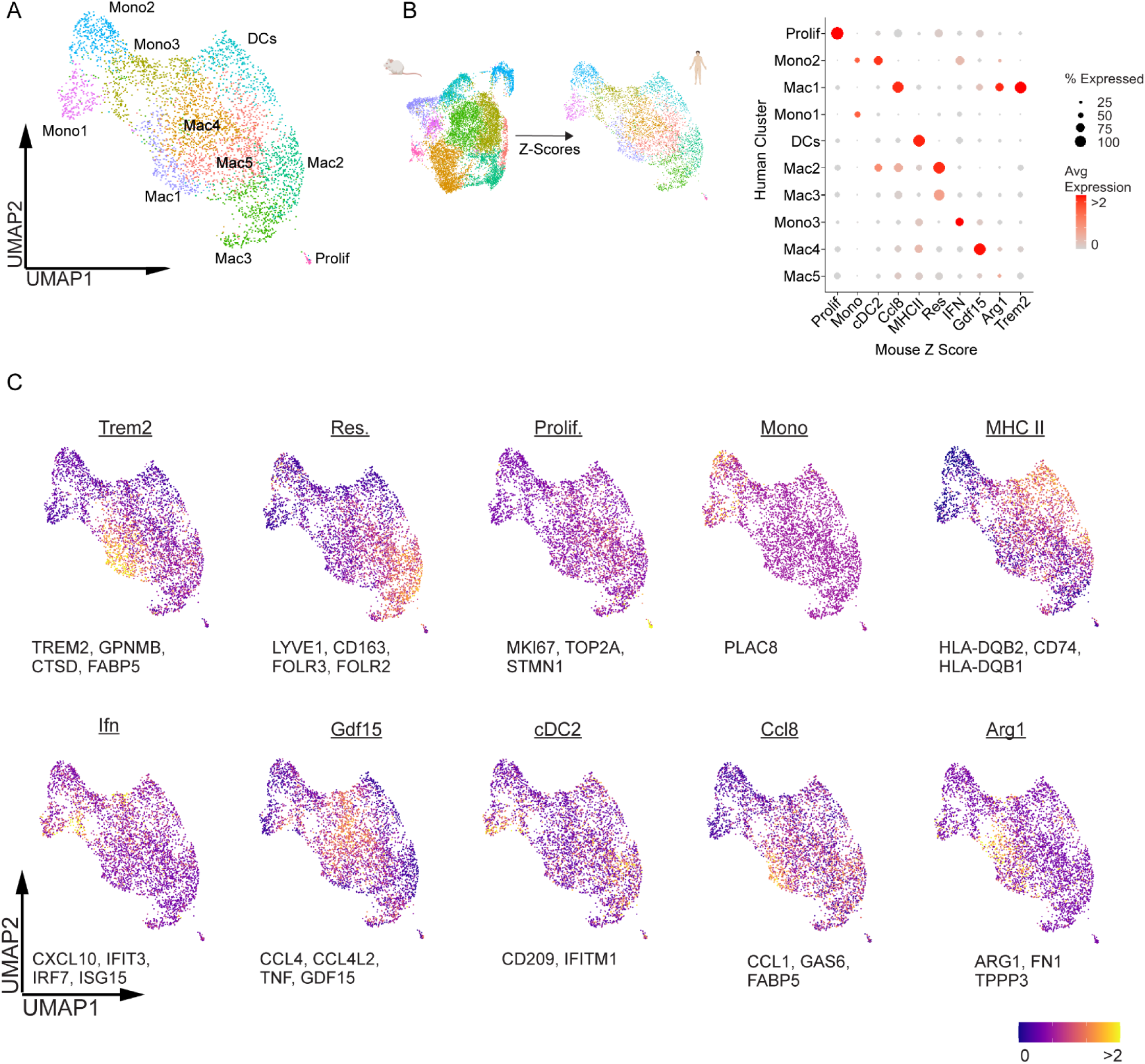
Monocyte-derived populations are conserved between mice and humans. **A,** UMAP plot of monocytes, macrophages, and dendritic cells from human donor (n=2) and heart failure samples (n=5). **B**, Schematic of analysis strategy (left) and dot plot of denoting the expression of genes enriched in each mouse monocyte, macrophage, and dendritic cell population (displayed as a z-score) in corresponding human monocyte, macrophage, and dendritic cell populations. Genes with mouse and human homologues are included in this analysis. **C**, UMAP feature plots of each mouse z-score signature in the human object.

### *Arg1*^+^ and interferon activated macrophages are predicted as early transient macrophage states

We utilized scVelo and Velocyto.R to perform RNA Velocity lineage tracing analysis (29, 30). We presented the data using t-distributed Stochastic Neighbor Embedding (t-SNE) projections split by time point post-MI (**Fig. 5A-B**). RNA Velocity suggested that monocytes differentiate into *Arg1^+^* and interferon activated macrophage populations as early intermediates, which subsequently differentiate to terminal *Trem2^+^* and MHCII^hi^ macrophages, respectively (**Fig. 5A**). Splitting the t-SNE plot by timepoint post-MI revealed two distinct populations of *Trem2^+^* macrophages, one arising at day 3 and a second evident beginning at day 7. The early *Trem2^+^* macrophage population expressed a distinct transcriptional signature that including *Arg1* and *Spp1*. The later *Trem2^+^* macrophage population displayed enriched expression of *Ninj1, Creg1,* and *Gstm1* (**Fig. 5A, Suppl. Fig. 4A).** Each of these *Trem2^+^* macrophage populations was also observed in a an independent trajectory analysis using Palantir (31). Palantir predicted that cDC2s, *Trem2*^+^ macrophages, and MHCII^hi^ macrophages were terminal cell states and that type-1 interferon activated macrophages and *Arg1^+^* macrophages represented intermediate states of differentiation based on entropy and pseudotime scores (**Fig. 5C)**. Plotting z-scores of each population also demonstrated increasing expression of terminal state markers further down the differentiation trajectories (**Fig. 5D**). Consistent with this framework, monocytes, *Arg1^+^* macrophages, and type-1 interferon activated macrophages were enriched at early time points post-MI and positioned directly adjacent to monocytes in pseudotime vs. entropy waterfall plots (**Suppl. Fig. 4B).**

**Fig. 5.**
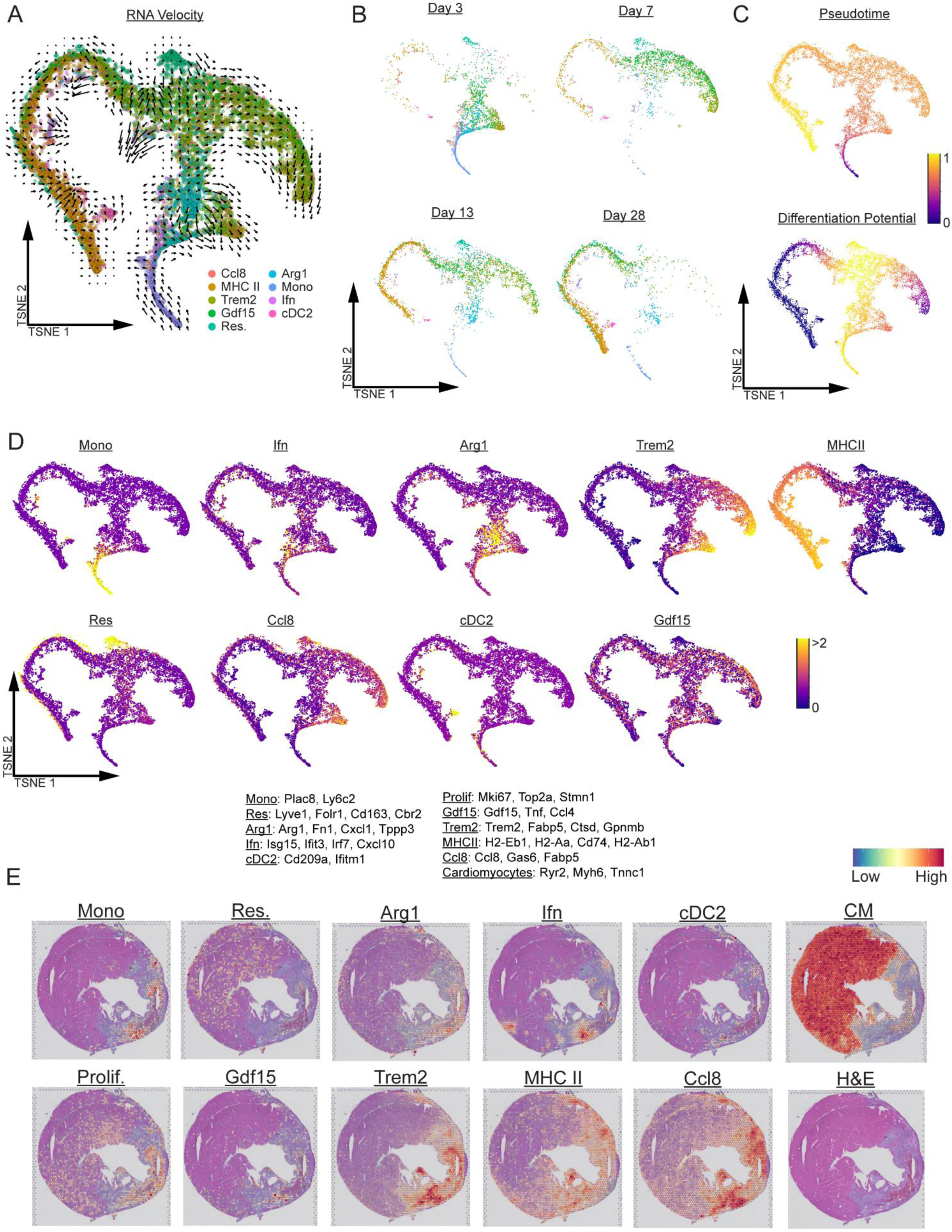
Monocyte-derived populations are spatially and temporally distinct. **A-B**, Palantir tSNE projection of combined time points labeled by cluster with superimposed RNA velocity information (A) and split by time point (B). **C**, Palantir tSNE projection labeled by pseudotime and differentiation potential. **D**, Feature plots of monocyte, macrophage and dependent cell populations z-scores overlayed on the tSNE projection. **E**, Spatial transcriptomic expression plots of monocyte, macrophage and dependent cell populations, and cardiomyocyte z-scores at day 7 post-MI.

To assess whether these diverse macrophage populations exist in different regions of the heart, we utilized a published spatial transcriptomic dataset of mouse MI.(32) We plotted z-scores of the individual macrophage states onto transverse heart sections at 7 days post-MI and observed that individual monocyte, cDC, and macrophage states were enriched in different regions of the infarct. The infarcted area can be visualized by loss of cardiomyocyte z-score expression*. Arg1^+^*, *Trem2^+^*, *Gdf15^+^*, and *Ccl8^+^* macrophages are enriched near the core of the infarct. Type-1 interferon activated macrophages, MHCII^hi^ macrophages, and cDC2s were enriched in the border zone. Resident macrophages are predominantly found outside of the infarct in the remote zone of the heart, while proliferating macrophages are distributed within the infarct and remote zones (**Fig. 5E**). Taken together, these data predict that monocytes differentiate into spatially restricted macrophage subsets and that type-1 interferon activated macrophages and *Arg1^+^* macrophages represent distinct intermediate macrophage populations.

### Recruited macrophages begin to acquire transcriptional identity during extravasation

To investigate where fate decisions are initially specified during the process of monocyte recruitment and extravasation into the heart, we performed closed chest ischemia-reperfusion injury on wild-type mice and harvested animals at 2 days post-MI. Mice were administered BV421 conjugated anti-CD45 antibodies retro-orbitally 5 minutes prior to harvest to label leukocytes in continuity with the circulation (intravascular compartment) (**Fig 6A**). Upon harvest, hearts were perfused to flush blood cells which had not adhered to the endothelium. Macrophages and monocytes were then sorted by FACS into intravascular (BV421+) and extravascular (BV421-) libraries for single cell RNA sequencing.

**Fig. 6.**
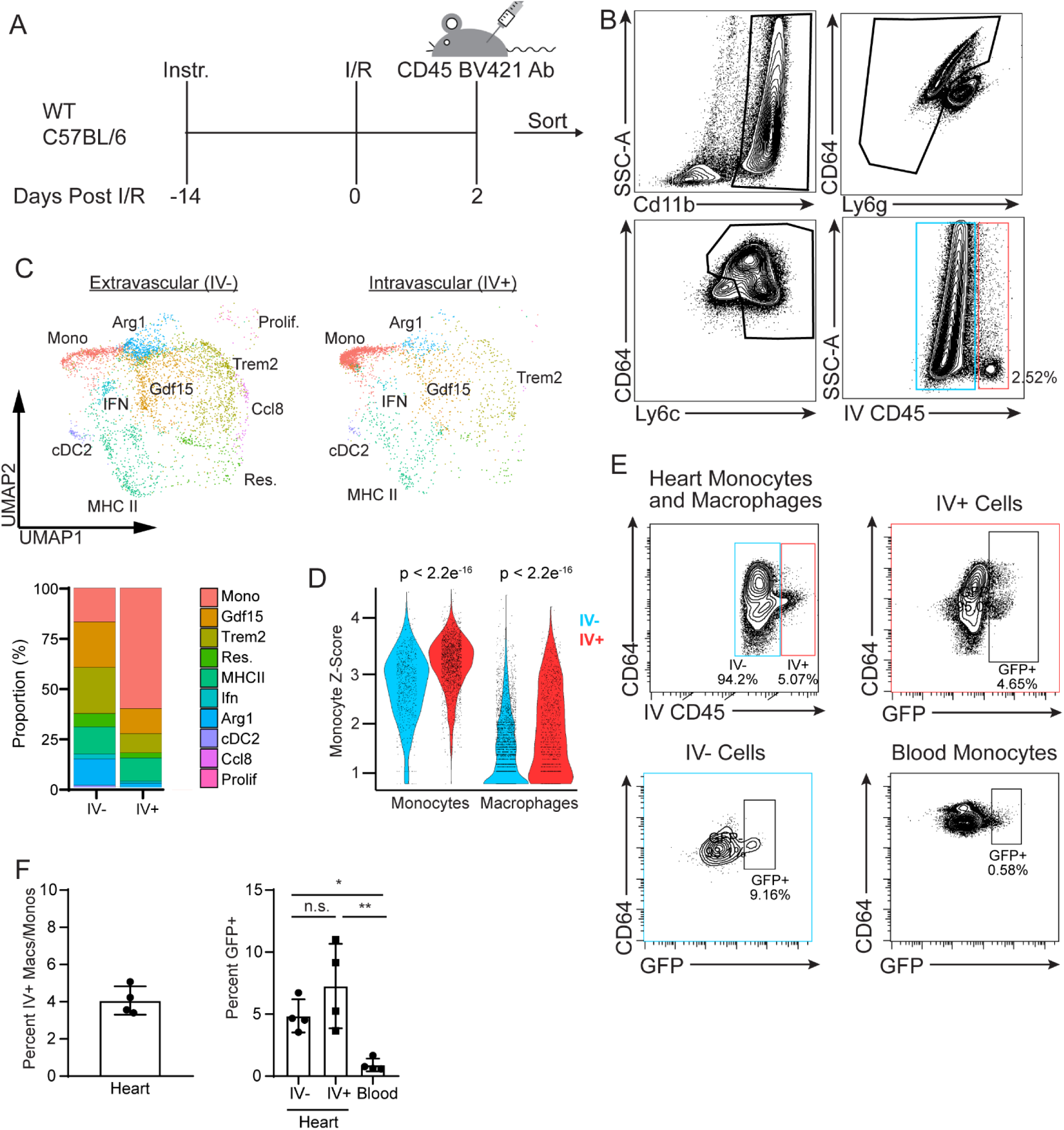
Monocyte fate specifications is initiated prior to myocardial infiltration. **A**, Experimental strategy for single cell RNA sequencing of intravascular (anti-CD45 BV421+) and extravascular (anti-CD45 BV421-) compartments. Anti-CD45 BV421 was administered via retro-orbital injection 5 minutes prior to collection. **B**, Representative FACS plots and gating strategy demonstrating anti-CD45 BV421 antibody staining in cardiac monocytes and macrophages. **C**, Clustered UMAP plots of intravascular (BV421+) and extravascular (BV421-) libraries and stacked bar chart showing distribution of monocyte, macrophage, and dendritic cell populations in each compartment. **D**, Violin plot of monocyte signature gene expression (expressed as a z-score) in monocytes and macrophages located within he intravascular and extravascular compartments. **E-F**, Representative FACS plots and quantification of GFP+ monocytes and macrophages within the heart and blood of *Mx1^GFP^* mice after MI. Intravascular and extravascular cardiac monocytes and macrophages expressing GFP is shown. * p<0.05, ** p<0.01

Approximately 2.5% of all macrophages and monocytes collected from the heart were BV421 positive, indicating that these cells had either adhered to the endothelium or were in the process of extravasating from the vasculature into the myocardium (**Fig. 6B, Suppl. Fig. 5**). Libraries were sequenced and cell clusters annotated by reference mapping to the *Ccr2^CreERT2^Rosa26^LSL-tdTomato^* monocyte lineage tracing object. As expected, the majority of intravascular cells mapped to the monocyte cluster. Interestingly, we also observe mapping of intravascular cells to identified macrophage populations suggesting that macrophage fate determination may begin during or prior to monocyte extravasation (**Fig. 6C**). Combined pseudotime analysis using the monocyte lineage tracing and IV anti-CD45 antibody labeled libraries also demonstrated distribution of intravascular cells throughout all clusters (**Suppl. Fig. 4C, D**). Consistent with reference mapping, marker genes enriched in each of the monocyte, macrophage, and dendritic cell clusters were expressed within corresponding cells in the both the intravascular and extravascular compartments (**Suppl. Fig. 5).** Monocytes and macrophages present within the intravascular compartment expressed transcriptional signatures of monocytes (monocyte z-score) suggesting that they recently differentiated from monocytes (**Fig. 6D**).

To corroborate observations from our sequencing data, we performed ischemia reperfusion injury on *Mx1^GFP^* mice (type I IFN reporter) and harvested hearts 2 days post-MI. Mice were injected with BV421 conjugated anti-CD45 antibodies 5 minutes prior to collection. FACS analysis revealed that 4% of monocytes and macrophages within the heart were BV421+ indicating that they were localized within the intravascular compartment. GFP positive cells were evident within the intravascular and extravascular monocyte and macrophage pool. Very few GFP expressing cells were detected in the blood. These findings confirm that type 1 interferon activation within monocytes and macrophages is initiated during extravasation (**Fig. 6E, F**). Taken together, these results demonstrate that cardiac infiltrating monocytes begin to acquire cell state specific transcriptional identity during extravasation and differentiation into macrophages.

### Type I interferon signaling in monocytes and monocyte-derived macrophages protects from adverse post-MI remodeling

To decipher the functional significance of these diverse macrophage populations, we focused on the IFN activated macrophage subset given their proximal position within the predicted differentiation trajectory and suggested clinical relevance (33). Utilizing the *Ccr2^CreERT2^Ifnar1^Flox^* mouse, we selectively deleted *Ifnar1* in recruited monocytes and their progeny using a single dose of tamoxifen injected IP (**Fig. 7A**).(34) Control and *Ccr2^CreERT2^Ifnar1^Flox^* mice were subjected to MI and echocardiography performed at day 28 post injury. *Ccr2^CreERT2^Ifnar1^Flox^* hearts demonstrated significantly decreased left ventricular ejection fraction, increased end systolic and diastolic left ventricular volumes, and increased akinetic area compared to controls (**Fig. 7B-C**). Analysis of cardiomyocyte size via wheat-germ agglutinin staining revealed increased cardiomyocyte hypertrophy in the border and remote zones of the left ventricle along with increased fibrosis measured by trichrome staining. We observed consistent findings in *Ccr2^CreERT2^Ifnar1^Flox^* mice subjected to continuous tamoxifen treatment, which will delete *Ifnar1* in resident CCR2+ macrophages, monocytes and their progeny (**Fig. 7D-F**). These data are consistent with the conclusions that type 1 interferon primarily functions within in monocytes and their progeny recruited to the heart early after injury and that type I IFN signaling within infiltrating monocytes and their progeny confers a protective effect on MI remodeling. Intriguingly, these findings are in contrast to studies demonstrating protective effects in whole body knockouts of *Ifnar1*, suggesting that type 1 interferon signaling has cell type specifics effects following (33).

**Fig. 7.**
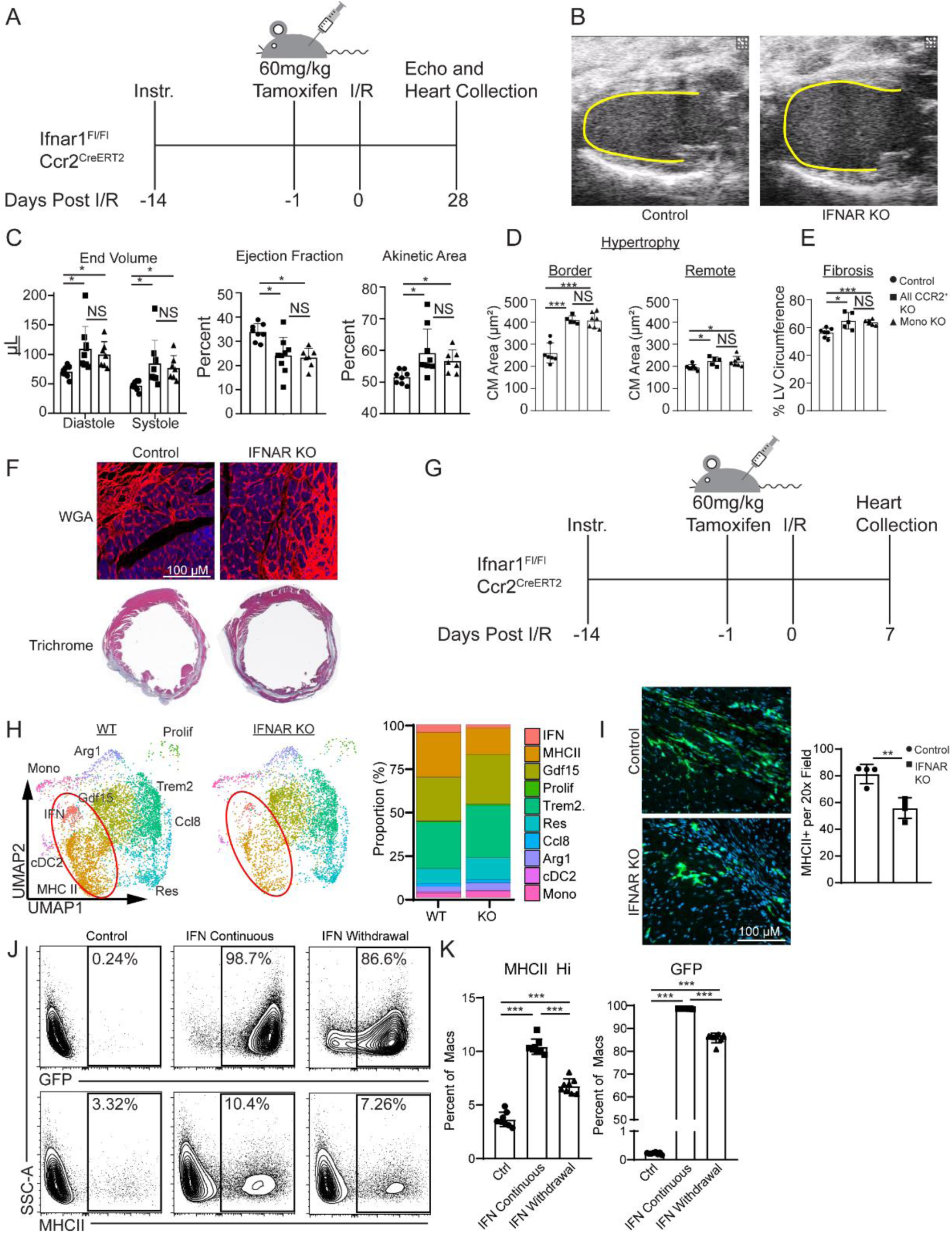
Type 1 interferon signaling in monocytes attenuate post-MI remodeling. **A**, Experimental strategy for deletion of *Ifnar* in recruited monocytes and their progeny. **B**, Representative long-axis echocardiographic images at end diastole showing enhanced LV remodeling in *Ccr2^ERT2Cre^Ifnar^Flox^* mice compared to controls. **C**, Quantification of echocardiography measurements of LV end systolic volume, ejection fraction, and akinetic area 4 weeks after MI. **D**, Measurement of cardiomyocyte size by WGA staining 4 weeks after MI. **E**, Measurement of infarct size assessed by fibrosis as a percentage of LV diameter 4 weeks after MI. **F**, Representative images of WGA and trichrome staining in control and *Ccr2^ERT2Cre^Ifnar^Flox^* hearts. **G**, Experimental strategy for single cell RNA sequencing of monocytes, macrophages, and dendritic cells in control and *Ccr2^ERT2Cre^Ifnar^Flox^* hearts. **H**, UMAP projection and stacked bar chart comparing distribution of cells in control and *Ccr2^ERT2Cre^Ifnar^Flox^* hearts. **I**, Representative images and bar chart quantifying MHCII+ macrophages per 20X field in control and *Ccr2^ERT2Cre^Ifnar^Flox^* hearts. **J-K**, FACS plots of elicited peritoneal macrophages isolated from Mx1^GFP^ mice stimulated with either PBS or *IFNβ* showing GFP and MHC-II expression (J). Quantification of the percent GFP+ and MHCII+ macrophages (K). * p<0.05, ** p<0.01, *** p<0.001

### Type I IFN signaling is essential for the differentiation of MHCII^hi^ macrophages

We next sought to determine the effect of *Ifnar1* knockout on the diversity of recruited monocytes and macrophages. We performed ischemia reperfusion injury on control and *Ccr2^CreERT2^Ifnar1^Flox^* mice given a single IP injection of tamoxifen 1 day prior to injury. Hearts were collected 7 days post injury and processed for single cell RNA sequencing (**Fig. 7G**). Libraries were sequenced and clusters annotated by reference mapping to the *Ccr2^CreERT2^Rosa26^LSL-tdTomato^* object. As expected, the proportion of interferon activated macrophages was reduced. Interestingly, the proportion of MHCII^hi^ macrophages was also dramatically reduced in *Ifnar1* knockout mice compared to littermate controls (**Fig. 7H, Suppl. Fig. 6A**). Combined pseudotime analysis using the monocyte labeling, IV labeling, and IFNAR KO libraries demonstrates a reduction in terminal MHCII^hi^ macrophages (**Suppl. Fig. 6B**). Immunostaining confirmed that MHCII^hi^ macrophages were significantly reduced in *Ifnar1* knockout hearts compared to controls at 28 days post injury (**Fig. 7I**).

To assess whether type I IFN signaling to monocytes directly influences MHCII^hi^ macrophage differentiation, we utilized thioglycolate induced peritoneal macrophages (pMacs) from *Mx1^GFP^* mice. pMacs were cultured in complete media with 25 nM IFNβ for 3 days followed by 4 days without IFNβ (IFN Withdrawal) or for 7 days with IFNβ (IFN Continuous). After 7 days of culture, pMacs were collected and flow cytometry performed to identify percentages of GFP+ and MHCII^hi^ macrophages. Continuous treatment with IFNβ resulted in GFP expression in 98.7% of pMacs compared to less than 0.25% of controls. Continuous treatment also significantly increased the proportion of MHCII^hi^ pMacs from 3.7% to 10.4%. Withdrawal of IFNβ reduced the percentage of MHC^II^ high macrophages showing a reversible effect (**Fig. 7J-K**). These results indicate that type I interferon signaling directly regulates the acquisition of the MHCII^hi^ macrophage fate.

## Discussion

This study provides new insights into the spatiotemporal dynamics of monocyte recruitment and differentiation following MI. Monocytes are recruited into the myocardium within the first 5-7 days post-MI and their progeny persisted through at least 28 days post-MI. Single cell RNA sequencing combined with genetic lineage tracing provided an unprecedented opportunity to precisely define the fates of monocytes that infiltrate the infarcted heart. This technique enabled the identification of monocyte, macrophage, and cDC2 populations that reside within the infarcted heart and demonstrated a coordinated and dynamic shift in macrophage phenotype over time. Furthermore, we demonstrated that these diverse macrophage subsets are situated in distinct locations within the heart, begin to acquire their transcriptional identity during extravasation, and that type I interferon activated macrophages and Arg1^+^ macrophages are predicted as intermediate populations that give rise to more terminal macrophage states. Within this framework, type I interferon signaling in infiltrating monocytes and derived macrophages promotes the differentiation of MHC-II^hi^ macrophages and confers protection following MI. These findings uncover the complexity of monocyte fate decisions and highlight the therapeutic potential for targeting monocyte differentiation.

In recent years, numerous publications have demonstrated the heterogeneity of macrophages in the heart and other organs using single cell RNA sequencing. Of note is the dramatically distinct transcriptional profiles taken on by recruited macrophages after tissue injury. However, little is known about when and how these distinct phenotypes are derived. To evaluate spatial localization of differentiation we leveraged a published spatial transcriptomics dataset of mouse MI which demonstrated distinct regions of differentiation relative to the infarcted area (32). Evaluating pseudotime identified intermediate IFN-activated and Arg1^+^ macrophage populations revealed these populations are concentrated in the border zone and core of the infarct, respectively. This finding is consistent with previous studies demonstrating clustering of the type 1 interferon response in the border zone.(35) These locations correspond to terminal population also found in these regions; MHCII^hi^ in the border zone and Trem2^+^ in the core, suggesting that localization within the infarct may influence the trajectory of recruited macrophages.

With respect to timing of fate acquisition, we observed that the majority of extravasating cells retained a monocyte expression profile. Retained expression of monocyte markers within intravascular macrophages is also consistent with the hypothesis that monocytes begin to express markers of specific macrophage populations during extravasation. Activation of the interferon reporter *MX1* in extravasating cells but not in the bone marrow further supports this hypothesis. A recent study using permanent ligation myocardial infarction in mice demonstrated interferon stimulated gene (ISG) expression in monocyte and neutrophil progenitors in the bone marrow as well as expression of ISGs in human peripheral blood neutrophils.(36) While our findings do not identify activation of interferon signaling in peripheral blood monocytes, they also do not rule out that activation within the bone marrow is possible in different injury models or using different detection methods. Future studies are needed to determine when during the extravasation process does macrophage fate specification begin.

This work lays the foundation for constructing monocyte differentiation trajectories. By analyzing single cell expression profiles at multiple timepoints after reperfused MI, we have demonstrated that recruited monocyte-derived macrophages are retained in tissue and alter their expression profile over time. Future studies are needed to direct trace the fates of each monocyte-derived subsets to defiantly understand whether they represent an intermediate or terminally differentiated state and understand their plasticity. The data presented here along with another recent publication (37), indicate that recruited monocytes differentiate into at least 2 intermediate states: *Arg1^+^* macrophages and type 1 interferon activated macrophages. We previously demonstrated through direct lineage tracing that *Arg1^+^* macrophages give rise to multiple additional macrophages subsets including *Gdf15^+^*, *Trem2^+^*, MHCII^hi^, and resident-like macrophages (37). Here, we demonstrate that interferon activated macrophages give rise to MHCII^hi^ macrophages. This is concept is supported by a dramatic decrease in MHCII^hi^ macrophages when *Ifnar1* is deleted in monocytes and derived macrophages and the observation that IFNβ treatment of elicited peritoneal macrophages *in vitro* is sufficient to induce expression of MHCII. Future experiments utilizing direct lineage tracing of interferon activated macrophages and other populations as well as unbiased lineage tracing techniques will further clarify trajectories of monocyte differentiation.

Previous studies have demonstrated that type 1 interferon signaling may have detrimental effects in the heart. MI induces a feed-forward type 1 interferon inflammatory response triggered by sensing of cytoplasmic DNA and activation of the cGAS-STING pathway in cardiomyocytes. Global deletion of STING, IRF3, and IFNAR in mice display reduce incidence of cardiac rupture, increased survival, and impaired wound healing after permanent left coronary artery ligation (33, 35). Similarly, pressure overload by transverse aortic constriction in mice also induces activation of type 1 interferon, reduced contractile function, and impaired sarcomeric turnover, phenotypes rescued by knockout of ISG15 (38). Conversely, our results demonstrate a cardioprotective effect of type 1 interferon signaling in monocytes and monocyte-derived macrophages, as evidenced by decreased left ventricular systolic function and accelerated cardiac remodeling after reperfused MI. Together, these data suggest that type 1 interferon signaling has cell specific functions in the injured and diseased heart and highlight the importance of targeted therapies to optimize the balance between risk and benefit.

This study is not without limitations. Computational analysis only predicts differentiation trajectories and relationships between each monocyte, macrophage, and dendritic cell population. Targeted and unbiased combinatorial genetic fate mapping is ultimately required to refine and validate our findings. The signaling pathways by which macrophage fates are specified within the vascular compartment remains obscure and will undoubtedly be the topic of future studies. Additionally, the precise mechanism by which interferon signaling orchestrates the differentiation of MHCII^hi^ macrophages and protects the heart after MI is of great interest and could lead to novel therapeutic insights.

In conclusion, we demonstrate that monocyte-derived macrophages are recruited to the heart within the first week following MI and differentiate into evolutionarily conserved, spatially restricted, and transcriptionally diverse array of distinct macrophage and dendritic cell-like states. We uncover that monocyte fate specification begins prior to entry into tissue and identify a novel role of type 1 interferon signaling in recruited monocytes and macrophages that governs the differentiation of MHCII^hi^ macrophages, preserves left ventricular systolic function, and prevents cardiac adverse remodeling. Together, our findings provide novel and clinically relevant insights that suggest monocyte fate specification may serve as a target of immunomodulatory therapy.

## Methods

### Sex as a biological variable

For all experiments involving mouse models, equal numbers of each sex were included in each experiment.

### Animal Models

Mus musculus were housed in ultraclean rodent barrier facility with veterinarians from the Washington University Division of Comparative Medicine on call or on site for animal care. All procedures were performed in accordance with IUCAC protocols. Strains used were *Cx3cr1^ERT2Cre^* (39) (JAX #020940), (JAX #007561), *Rosa26^lox-stop-lox-tdTomato^* (40) (JAX #007914), *CCR2^ERT2Cre^* (41), *MX1^GFP^* (42) (JAC #033219), and *Ifnar1^Flox^* (34) (JAX # 028256) on the C57BL/6 background. Experiments were performed on mice 6-12 weeks of age. Equal numbers of male and female mice were used for experiments. For Cre recombination in all monocytes and macrophages, *Ccr2^CreERT2^ Ifnar1^flox/flox^* mice were given tamoxifen food pellets (Envigo TD.130857) for the entirety of the experiment. For Cre recombination in other lines, mice were given IP injection of 60 mg/kg of tamoxifen (Millipore Sigma T5648) at the indicated frequency and timing.

### Closed-chest Ischemia Reperfusion Injury

Mice were anesthetized, intubated, and mechanically ventilated. The heart was exposed, and a suture placed around the proximal left coronary artery. The suture was threaded through a 1 mm piece of polyethylene tubing to serve as the arterial occlude. Each end of the suture was exteriorized through the thorax. The skin was closed, and mice were given a 2-week recovery period prior to induction of ischemia. After 2 weeks, the animals were anesthetized and placed on an Indus Mouse Surgical Monitor system to accurately record ECG during ischemia. Ischemia was induced after anesthetizing the animals. Tension was exerted on suture ends until ST-segment elevation was seen via ECG. Following 90 minutes of ischemia time, tension was released, and the skin was then closed.

### Echocardiography

Mice were sedated with Avertin (0.005 ml/g) and 2D and M-mode images were obtained in the long and short axis views 4 weeks after MI using the VisualSonics770 Echocardiography System. Left diastolic volume, left systolic volume, akinetic area, and total LV area was measured using edge detection and tracking software (VivoLab). Ejection fraction was calculated as the (left diastolic volume – left systolic volume) / left diastolic volume. Akinetic percentage was calculated as the akinetic area / total LV area.

### Histology

4 weeks after MI, mice were euthanized, and hearts perfused with PBS. Hearts were fixed overnight with 4% PFA in PBS at 4°C, cut into thirds on the transverse plane, placed into histology cassettes for paraffin embedding, and dehydrated in 70% ethanol in water. Hearts were paraffin embedded and cut into 4 µm sections and mounted on positively charged slides. Tissues were stained by Gomori’s Trichrome staining (Richard-Allan Scientific 87020) and WGA staining (Vector Laboratories RL-1022). Trichrome staining was imaged using a Meyer PathScan Enabler IV. Measurement of infarct length and LV length was performed in FIJI, with data quantification normalizing the length of the infarct to the length of the LV(43). WGA staining was imaged using the 20x objective of a Zeiss Axioscan7 confocal microscope. 3 20x fields of the border zone of the infarct were randomly selected per mouse. 20 cross sectional (not longitudinal) cardiomyocytes per 20x field were selected at random. Measurement of cross-sectional cardiomyocyte area was performed in FIJI. Data is displayed as the average cardiomyocyte cross sectional area per mouse.

### Immunofluorescence

Mice were euthanized, and hearts perfused with PBS. Hearts were cut midway on the transverse plane and fixed overnight with 4% PFA in PBS at 4°C. They were then dehydrated in 30% sucrose in PBS overnight at 4°C. Hearts were embedded in O.C.T. (Sakura 4583) and frozen at −80°C for 30 minutes. Hearts were then sectioned at 15 µm using a Leica Cryostat, mounted on positively charged slides, and stored at −80°C. For staining, the following steps were performed at room temperature and protected from flight. Slides were brought to room temperature for 5 minutes and washed in PBS for 5 minutes. Sections were then permeabilized in 0.25% Triton X in PBS for 5 minutes. Sections were then blocked in 5% BSA in PBS for 1 hour. Sections were then stained with primary antibody diluted in 1% BSA in PBS for 1 hour with rat anti CD68 (BioLegend 137002 1:400 dilution), rabbit anti Lyve1 (abcam ab14917 1:200 dilution). Sections were then washed 3 times 5 minutes each with PBS-Tween. After washing, the appropriate secondary antibodies were added diluted in 1% BSA in PBS for 1 hour at a 1:1000 dilution (Goat anti Rat 488 Life Technologies a11006, Donkey anti Rabbit 647 Abcam ab150075,). Slides were again washed 3 times 5 minutes each with PBS-T, and then mounted with DAPI mounting media (Millipore Sigma F6057) and cover slipped. Slides were stored at 4°C protected from light and imaged within 1 week.

Slides were imaged using the 20x objective of a Zeiss Axioscan7 microscope. Visualization of tdTomato was from the endogenous signal. For quantification, 3-5 20x fields were randomly selected per mouse of either the remote zone or infarct zone as indicated. Quantification of cell numbers were performed in FIJI or Zeiss Zen, and data is displayed as the average of each 20x field per mouse.

### Flow Cytometry

Hearts were perfused with PBS, weighed, minced, and digested for 45 minutes at 37°C in DMEM (Gibco 11965-084) containing 4500 U/ml Collagenase IV (Millipore Sigma C5138), 2400 U/ml Hyaluronidase I (Millipore Sigma H3506), and 6000 U/ml DNAse I (Millipore Sigma D4527). Enzymes were deactivated with HBSS containing 2% FBS and 0.2% BSA and filtered through 40-micron strainers. Cells were incubated with ACK lysis buffer (Gibco A10492-01) for 5 minutes at room temperature. Cells were washed with DMEM and resuspended in 100 µl of FACS buffer (PBS containing 2% FBS and 2mM EDTA).

Approximately 100 μl of blood was collected by cheek bleed into a tube containing 20 μl of EDTA (Corning 46-034-CI). Cells were incubated with ACK lysis buffer for 15 minutes at room temperature, washed, and ACK lysed a second time. Cells were then washed with DMEM and resuspended in 100 μl of FACS buffer.

Cells were incubated in a 1:200 dilution of fluorescence conjugated monoclonal antibodies for 30 minutes at 4°C. Samples were washed in FACS buffer and resuspended in a final volume of 250 µl FACS buffer. Flow cytometry and sorting was performed using a BD FACS melody.

For sorting of monocytes and macrophages from *Cx3cr1^CreERT2^Rosa26^LSL-tdTomato^* for single cell RNA sequencing, antibodies used were: CD45 PerCP/Cy5.5 (BioLegend 103132), Ly6g APC/Cy7 (BioLegend 127623), Ly6c APC (BioLegend 128016), CD64 PE/Cy7 (BioLegend 139313), and Cd11b BV421 (BioLegend 101235).

For sorting of monocytes and macrophages from *Ccr2^CreERT2^Rosa26^LSL-tdTomato^* for single cell RNA sequencing, antibodies used were: CD45 PerCP/Cy5.5 (BioLegend 103132), Ly6g APC/Cy7 (BioLegend 127623), Ly6c FITC (BioLegend 128005), and CD64 APC (BioLegend 164409), Cd11b PE/Cy7 (BioLegend 101215), CD3 BV510 (BioLegend 100233), CD19 BV510 (BioLegend 115545) and MHCII BV421 (BioLegend 107631).

For flow cytometry of blood from *Ccr2^CreERT2^Rosa26^LSL-tdTomato^*, antibodies used were: CD45 PerCP/Cy5.5 (BioLegend 103132), Ly6g BV421 (BioLegend 127627), Ly6c PE/Cy7 (BioLegend 128017), CD115 APC (BioLegend 135509), CD3 FITC (BioLegend 152303), and CD19 BV510 (BioLegend 115545).

For flow cytometry of bone marrow from *Ccr2^CreERT2^Rosa26^LSL-tdTomato^*, antibodies used were: Sca1 FITC (BioLegend 108105), c-Kit PerCP/Cy5.5 (BioLegend 135133), Il7ra PE/Cy7 (BioLegend 158209), CD34 APC (BioLegend 119309), CD16/CD32 APC/Cy7 (BioLegend 156611), and Lineage Cocktail BV421 (BioLegend 133311).

### Single Cell RNA Sequencing Analysis

Monocytes and macrophages were sorted after I/R as indicated. Single cell RNA sequencing libraries were generated using the 10X Genomics 5’v1 platform. Libraries were sequenced using a NovaSeq 6000 at the McDonnel Genome Institute (MGI). For *Cx3cr1^CreERT2^Rosa26^LSL-tdTomato^* mice, cells were separated into tomato- and tomato+ libraries. Sequencing reads were aligned to the GRCm38 transcriptome using CellRanger (v3.1.0) from 10X Genomics. Custom

Downstream data processing was performed in the R package Seurat (v4)(44). SCtransform was used to scale and normalize the data for RNA counts. Quality control filters of genes/cell>200, mitochondrial reads/cell<10%, read counts/cell>5000, and read counts/cell<40000 were applied. PCA was used for dimensionality reduction, data was clustered at multiple resolutions, and UMAP projections were generated for data visualization. Harmony integration was applied to account for batch effect from different time points(45). Contaminating populations of neutrophils and non-myeloid cells were excluded from analysis. Differential gene expression of upregulated genes in each subpopulation was generated in Seurat using FindAllMarkers using the Wilcoxan Rank Sum Test with a minimum fraction (min.pct) of 0.1 and a logFC threshold of 0.1. Gaussian kernel density estimation plots were generated using the Python package Scanpy (v1.8.1)(46).

### RNA Velocity Analysis

The ScVelo Python package was used to generate loom files with spliced and unspliced read counts for each library (29). Velocyto.R package for R was then utilized to perform RNA velocity analysis using the TSNE generated by Palantir (30). RunVelocity command was performed using default parameters of spliced.average = 0.2 and unspliced.average = 0.05. Velocity is plotted on the TSNE using the show.velocity.on.embedding.cor command using a neighborhood size of 200 cells and square-root scaling of velocity.

### Palantir Trajectory Analysis

The Palantir Python package was used for trajectory analysis (v1.0.0 to v1.1)(31). Prior to inputting data to Palantir, cell cycle regression was performed using Seurat and proliferating cells were excluded from pseudotime analysis. Monocytes were used as the early and starting cell type for Palantir. Highly variable gene analysis was not used. The eigenvectors used was determined by the eigengap. The number of waypoints used was 500. The option to use the early cell as the start cell was used. The output pseudotime/entropy values, terminal state probabilities, and TSNE coordinates for each cell were used to generate plots in R.

### Spatial transcriptomics

Spatial transcriptomics data of macrophage sub-population localization was generated using a published data set of mouse MI 7 days after injury (GSE176092)(32). Spatial data was analyzed using Seurat. The macrophage population gene signature was plotted by calculating the Z-score of the genes as listed and using the function “SpatialFeaturePlot” using the Z-score as a feature.

### Statistical Analysis

For analysis of echocardiography, histology, and immunofluorescence, two tailed t-test with Welch’s correction was performed in GraphPad Prism. At least 3 replicates were used for experiments. For percentage of tdTomato positive cells by FACS and immunofluorescence, a Brown-Forsythe one-way ANOVA was perfromed in Prism. P < 0.05 was a priori considered statistically significant. Data was plotted using Prism software.

### Study Approval

Animal studies were performed in compliance with guidelines set forth by the National Institutes of Health Office of Laboratory Animal Welfare and approved by the Washington University institutional animal care and use committee.

## Data Availability

Single Cell RNA sequencing data is available on the Gene Expression Omnibus (GSE252086).

## Code Availability

R and Python scripts are available on GitHub (https://github.com/alkoenig/Cardiac_Monocyte_Fate).

## Author Contributions

A.L.K. generated single-cell sequencing libraries. A.L.K, J.A, and F.F. performed the analysis of single-cell and single-nucleus sequencing data. A.L.K., G.S., and S.Y. performed histology and immunofluorescence experiments. K.J.W., J.M.N., and A.K. of the Washington University Mouse Cardiovascular Phenotyping Core performed myocardial infarction and echocardiography procedures. K.L. and A.L.K. are responsible for all aspects of this manuscript including experimental design, data analysis and manuscript production.

## Acknowledgements

ALK is supported by funding provided by the National Institutes of Health (K99HL166861) and the Myocarditis Foundation postdoctoral award (P22-04488). KJL is supported by the Washington University in St. Louis Rheumatic Diseases Research Resource-Based Center grant (NIH P30AR073752), the National Institutes of Health [R01 HL138466, R01 HL139714, R01 HL151078, R01 HL161185, R35 HL161185], Leducq Foundation Network (#20CVD02), Burroughs Welcome Fund (1014782), and Children’s Discovery Institute of Washington University and St. Louis Children’s Hospital (CH-II-2015-462, CH-II-2017-628, PM-LI-2019-829), Foundation of Barnes-Jewish Hospital (8038-88), and generous gifts from Washington University School of Medicine. We are thankful to the Mouse Cardiovascular Phenotyping Core facility at Washington University for performing mouse echocardiography and the reperfused myocardial infarction surgeries. We are thankful to the Genome Technology Access Center at the McDonnell Genome Institute for help with genomic analysis, the Digestive Diseases Research Core Center (DDRCC) for histology services and the Washington University Center for Cellular Imaging (WUCCI) at Washington University School of Medicine.

