## Supplementary material for "Genetic Mapping of Monocyte Fate Decisions Following Myocardial Infarction": Supplmental Figures

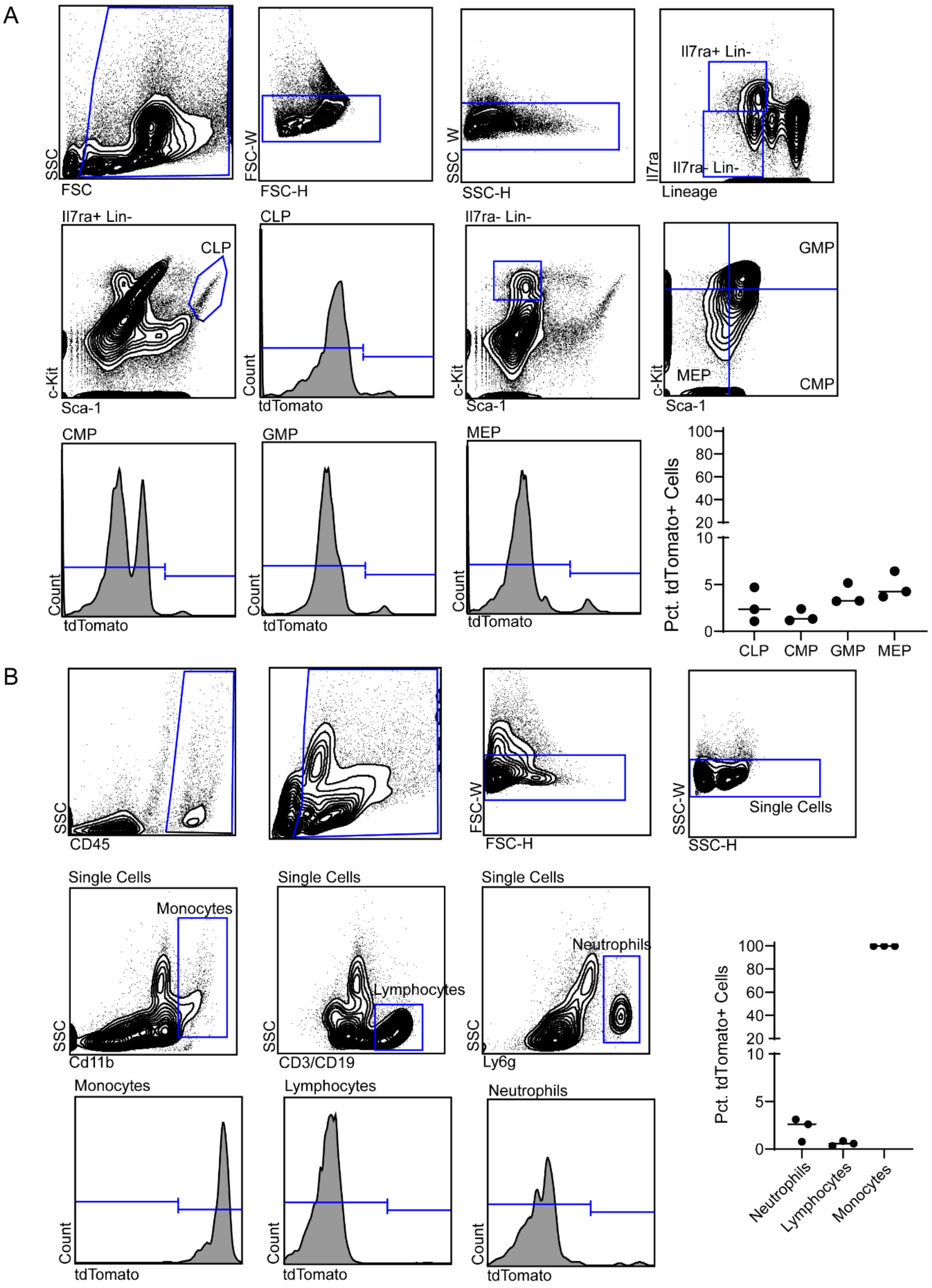


**Suppl. Fig. 1 Bone marrow and blood FACS gating strategies. A,** Representative gates and quantification of the percent of tdTomato positive progenitors cells in the bone marrow by flow cytometry in *Ccr2*$^{CreERT2}$*Rosa26*$^{LSL\text{-}tdTomato}$ mouse 1 day after tamoxifen injection. CLP: common lymphoid progenitor, CMP: common myeloid progenitor, GMP: granulocyte-monocyte progenitor, MEP: megakaryocyte-erythrocyte progenitor). **B**, Representative gates and quantification of the percent of tdTomato positive cells in the blood by flow cytometry in *Ccr2*$^{CreERT2}$*Rosa26*$^{LSL\text{-}tdTomato}$ mouse 1 day after tamoxifen.

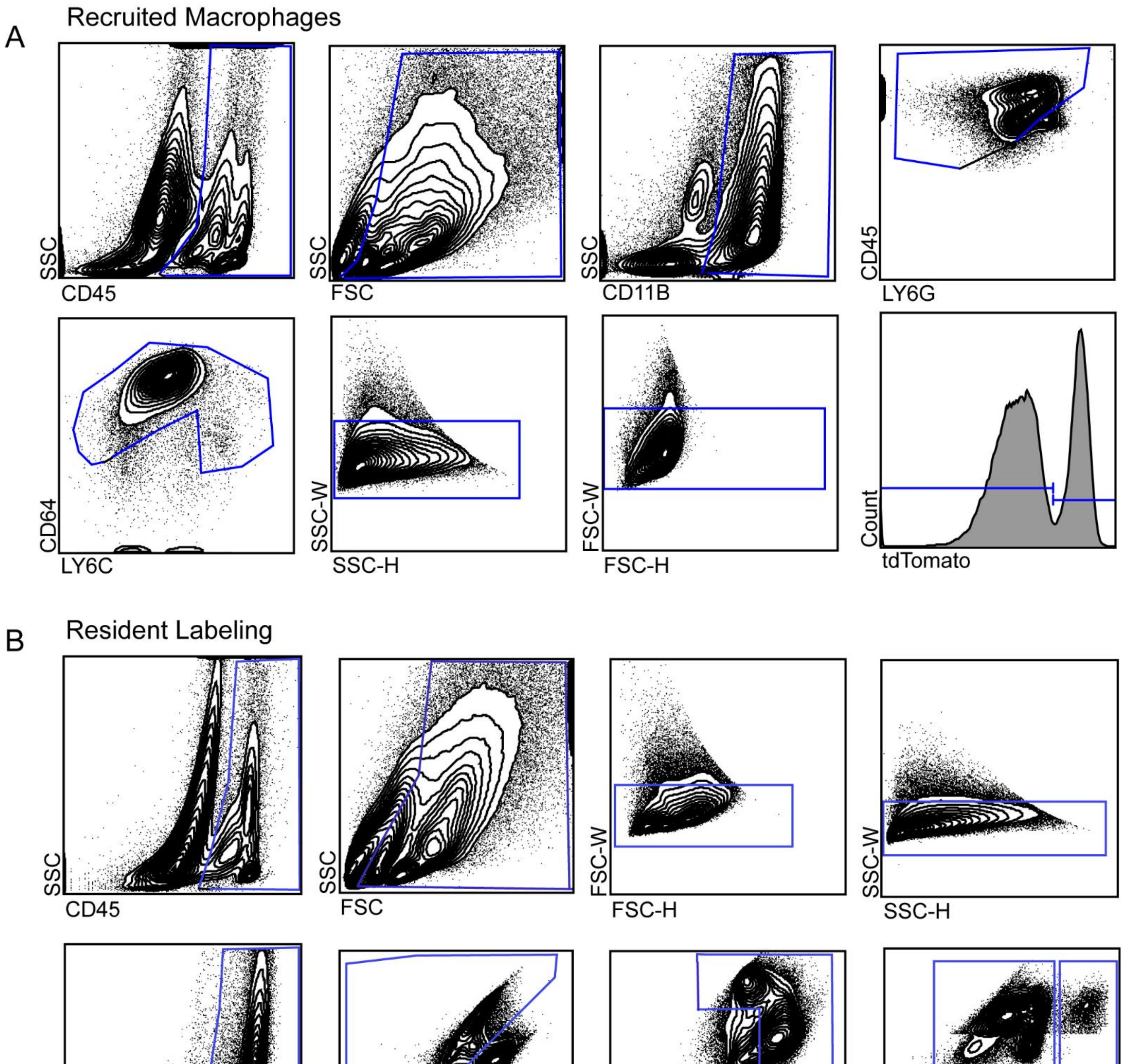


**Suppl. Fig. 2 FACS gating strategies for constructing monocyte and cardiac resident macrophage single cell RNA sequencing libraries. A**, Flow cytometry gating strategies used to isolate cardiac monocytes and macrophages for monocyte lineage tracing single cell RNA sequencing libraries. **B**, Flow cytometry gating strategies used to isolate cardiac monocytes and macrophages for cardiac resident macrophage lineage tracing single cell RNA sequencing libraries.

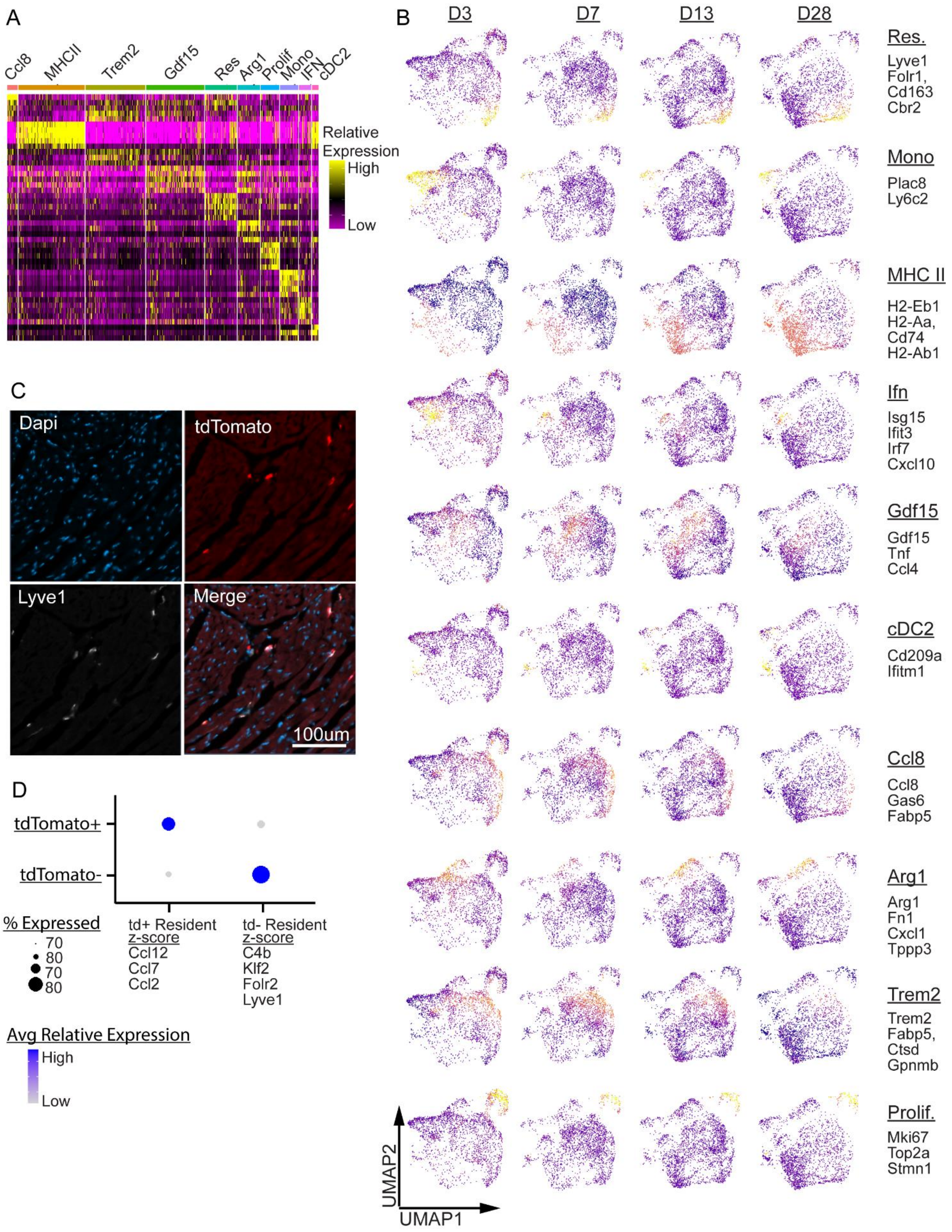


**Suppl. Fig. 3 Single cell RNA sequencing of recruited monocytes and their progeny after MI. A**, Heatmap of top 5 enriched genes in each monocyte, macrophage, and dendritic cell cluster. **B**, UMAP projections of marker genes (expressed as z-scores) for each cluster split by time point post-MI. **C**, Representative immunostaining image of LYVE1 expression in tdTomato+ macrophage in *Ccr2$^{CreERT2}$Rosa26$^{lsl-tdTomato}$* hearts 4 weeks after MI. **D**, Dot plot of transcriptional signatures that differentiate tdTomato- resident cardiac macrophages and tdTomato+ resident-like cardiac macrophages.

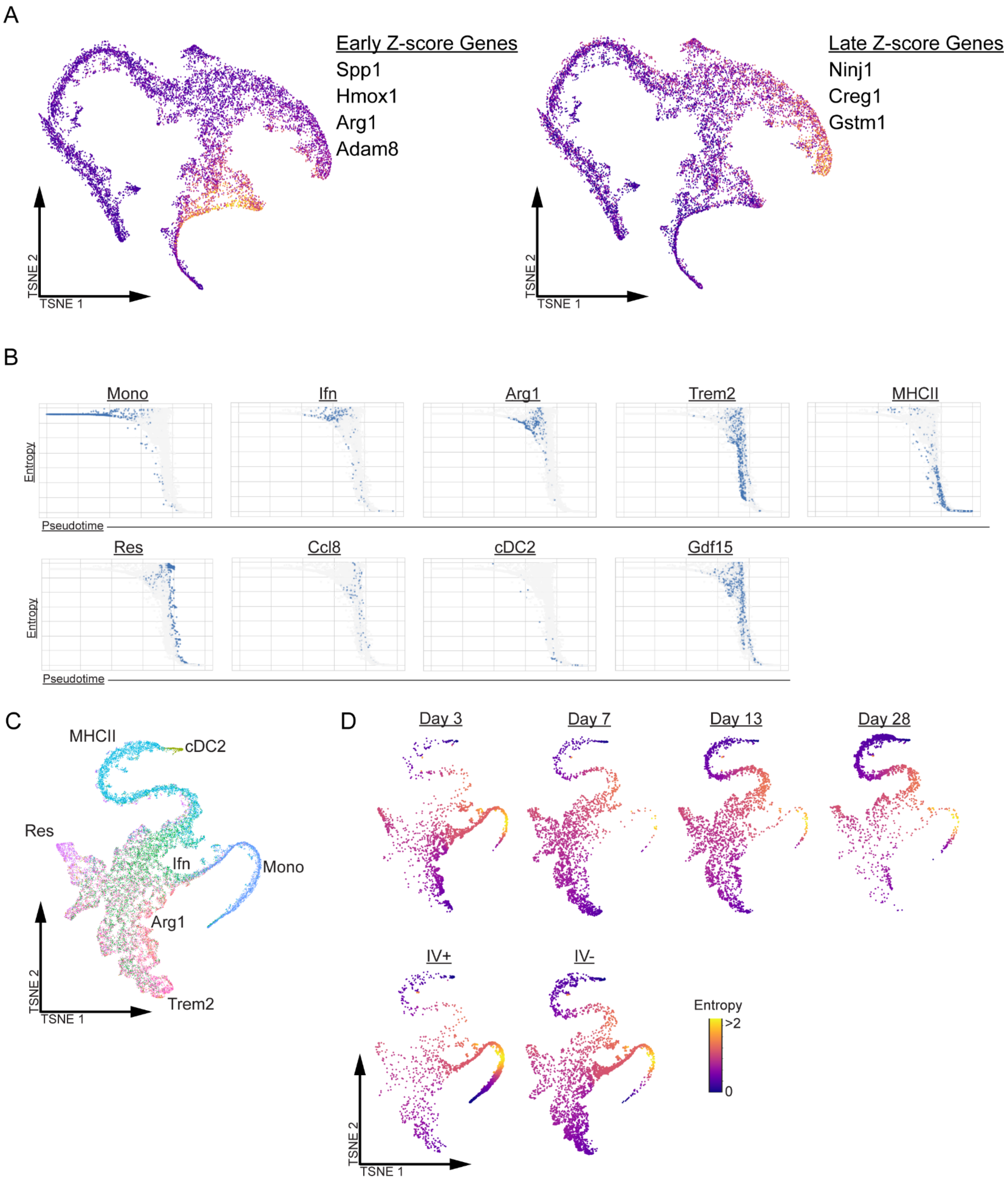


**Suppl. Fig. 4 Monocyte differentiation trajectories predicted by pseudotime analysis. A**, tSNE feature plot of transcriptional signatures (displayed as a z-score) enriched in the early (left) and late (right) *Trem2+* macrophage populations. **B**, Waterfall plots of pseudotime versus differentiation potential split by cluster. **C**, tSNE plots split by condition (lineage tracing and anti-CD45 antibody IV labeling) showing entropy values.

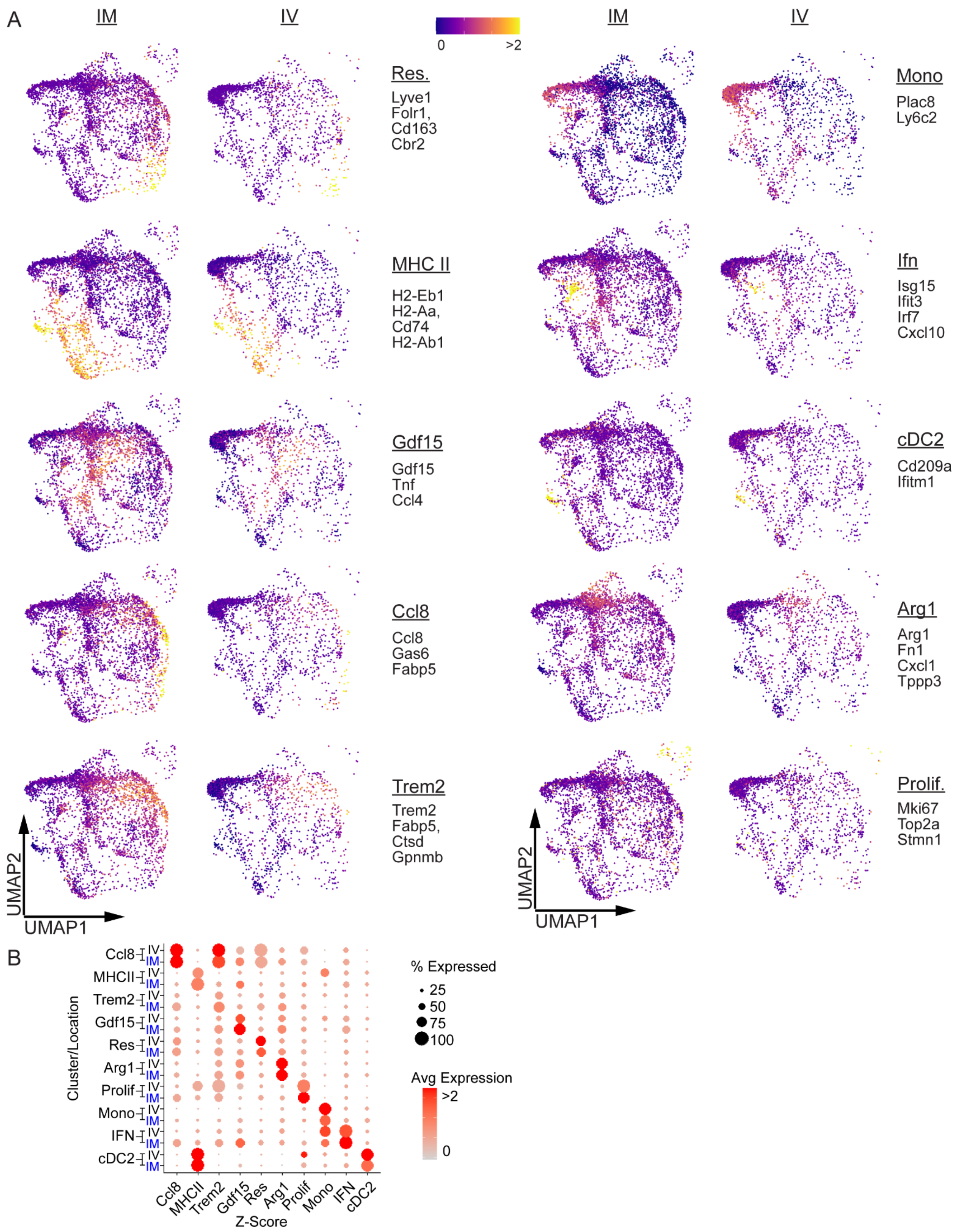


**Suppl. Fig. 5 Single cell RNA sequencing of intravascular and extravascular monocytes and macrophages after MI. A**, UMAP projections of genes (displayed as z-scores) enriched in each cluster split by intravascular vs. extravascular compartment. **B**, Dot plots of marker gene z-scores in each cluster split by intravascular vs. extravascular compartment.

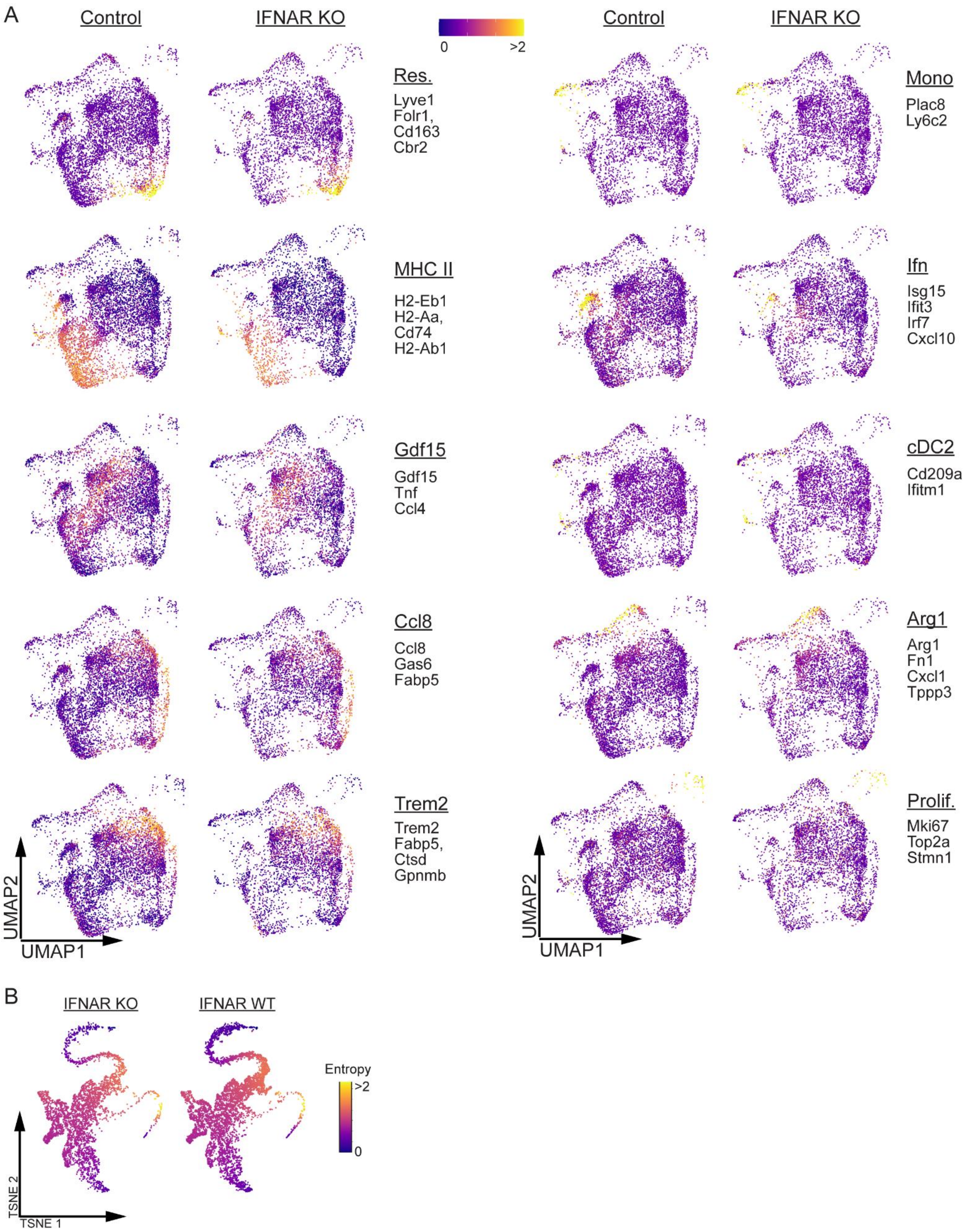


**Suppl. Fig. 6 Single cell RNA sequencing of monocytes and macrophages from control and *Ccr2*$^{ERT2Cre}$*Ifnar*$^{Flox}$ hearts. A**, UMAP projections of genes enriched (displayed as z-scores) in each cluster split by experimental condition. **B**, tSNE plots showing entropy values split by experimental condition.
